# Metabolic Control of Viral Infection through PPAR-α Regulation of STING Signaling

**DOI:** 10.1101/731208

**Authors:** Lili Tao, Alexandria Lowe, Guoxun Wang, Igor Dozmorov, Tyron Chang, Nan Yan, Tiffany A. Reese

## Abstract

Peroxisomal proliferator activated receptors (PPARs) are sensors of dietary lipids and pharmacological targets in the treatment of metabolic disorders. PPAR ligands are also immunosuppressive. However, their function during infection is debated and the mechanisms that underlie their immunoregulatory properties are unclear. We investigated the consequences of PPAR activation during herpesvirus infection. We found that activation of PPAR-α increased herpesvirus replication, suppressed type I interferon production and induced reactive oxygen species (ROS). We discovered that ROS induced by PPAR-α stimulation suppressed the cytoplasmic DNA sensing pathway after direct activation of stimulator of interferon (STING), the ER adapter downstream of cytoplasmic DNA recognition. Although high ROS induces inflammasome activation and cytokine production, we found that ROS inhibited interferon production after cytoplasmic DNA recognition. Treatment of mice with a clinically relevant agonist of PPAR-α increased herpesvirus replication and pathogenesis, comparable to levels observed previously in type I interferon receptor knockout mice. These findings reveal that activation of PPAR-α regulates immunity to cytoplasmic DNA and DNA virus infection through inhibition of interferon. Moreover, these results demonstrate that STING signaling and interferon production is regulated by ROS.

## Introduction

Peroxisome proliferator activated receptors (PPARs) are ligand activated nuclear receptors that regulate fatty acid oxidation and cholesterol metabolism. The PPAR family consists of three isoforms: PPAR-α, PPAR-β/δ, and PPAR-γ. All three receptors regulate lipid homeostasis, although they differ in terms of their tissue distribution, ligand specificities, and gene targets^1^. There are many natural and synthetic ligands of PPARs that are used to treat dyslipidemia and glucose disorders. Natural ligands include dietary lipids, such as omega-3 polyunsaturated fatty acids found in fish oil, and immune mediators, such as eicosanoids. Synthetic ligands, including fibrates that target PPAR-α and thiazolidinediones that target PPAR-γ, are used to treat metabolic disorders^1,2^.

In addition to their role in metabolism, PPARs regulate inflammation. PPAR agonists reduce inflammation in atherosclerosis, diabetes, neurodegenerative diseases, and autoimmune diseases^3^. Their mechanisms of action are diverse and include repression of NFκB and AP-1 DNA binding, regulation of nitric oxide, inhibition of dendritic cell maturation, reduction of cytokine expression by effector T cells, and inhibition of leukocyte recruitment to sites of inflammation^4,5^. Despite their immunosuppressive function, the effects of PPAR agonists on infectious disease outcomes has not been extensively studied. There are contradictory reports that synthetic agonists or dietary lipids improve or impair resistance to pathogen challenge, but we do not know the molecular mechanisms of PPAR-mediate immunoregulation during infection^6–10^.

Intermediates of cellular metabolism are important signaling molecules that alter immune defense pathways^11^. PPARs, as their name implies, increase peroxisomal metabolism. Peroxisomes synthesize phospholipids and bile acids and oxidize very long, branched chain, and polyunsaturated fatty acids. They also produce significant amounts of hydrogen peroxide and other reactive oxygen species (ROS). ROS have been widely implicated in inflammatory processes, particularly for NFκB signaling and inflammasome activation^12^. In general, antioxidant treatment reduces inflammatory cytokines and interferon-β (IFNβ) after stimulation with toll-like receptor agonists or viral RNA^13–16^. However, it is unclear whether peroxisomal metabolism and ROS function in antiviral immunity to DNA viruses and cytoplasmic DNA.

Herpesviruses manipulate metabolism during infection to promote viral replication and chronic infection^17,18^. In addition to altering glucose metabolism, herpesviruses induce peroxisome proliferation^19,20^. Moreover, they encode viral proteins that target peroxisomes, suggesting that modulation of peroxisomal function is important for these viruses^21–23^.

Here we investigated the consequences of activation of PPARs and peroxisomal metabolism on herpesvirus infection and induction of antiviral immune responses. We compared agonists for the three different PPARs and found that activation of PPAR-α increased herpesvirus replication. PPAR-α stimulation inhibited type I interferon production following DNA-virus infection or direct stimulation of the cyclic GMP-AMP synthase (cGAS)/stimulator of interferon (STING) pathway. Importantly, treatment of mice with PPAR-α agonist led to increased herpesvirus replication and heightened lethality in agonist-treated mice. Our data suggest that activation of PPAR-α suppresses cytoplasmic DNA sensing by generating high ROS that inhibit STING activation, leading to reduced type I interferon and impaired immunity to viral infections.

## Results

### Activation of PPAR-α promotes MHV68 replication

To examine the effects of PPAR activation on DNA virus infection, we used murine gammaherpesvirus-68 (MHV68) as our model. MHV68 readily infects mice and undergoes phases of infections similar to human herpesvirus, such as Kaposi’s sarcoma associated herpesvirus (KSHV) and Epstein Barr virus (EBV)^24–26^. We treated bone marrow derived macrophages with PPAR-α agonists, fenofibrate or WY14643, PPAR-β/δ agonist GW501516, or PPAR-γ agonist rosiglitazone. Following pretreatment, cells were infected with MHV68. We examined expression of lytic viral proteins on infected cells by flow cytometry to assay for viral replication^25^. We found that PPAR-α agonists fenofibrate and WY14643 both increased expression of lytic viral proteins on infected macrophages compared with untreated cells or cells treated with GW501516 or rosiglitazone (Figure 1a). MHV68 replication at either low or high multiplicity of infection (MOI) was also increased in macrophages treated with fenofibrate or WY14643 compared with untreated cells (Figure 1b, c, Supplemental Figure 1). Treatment with agonist had no effect on virus replication in macrophages isolated from *Ppara^−/−^* mice, suggesting that PPAR-α is critical for agonist effects (Figure 1d). We tested whether stimulation of PPAR-α affected replication of another herpesvirus that also infects macrophages and found that replication of murine cytomegalovirus (MCMV) was also increased by PPAR-α agonist (Supplemental Figure 2). Thus, PPAR-α stimulation increases herpesvirus replication in a PPAR-α-dependent manner.

**Figure 1.**
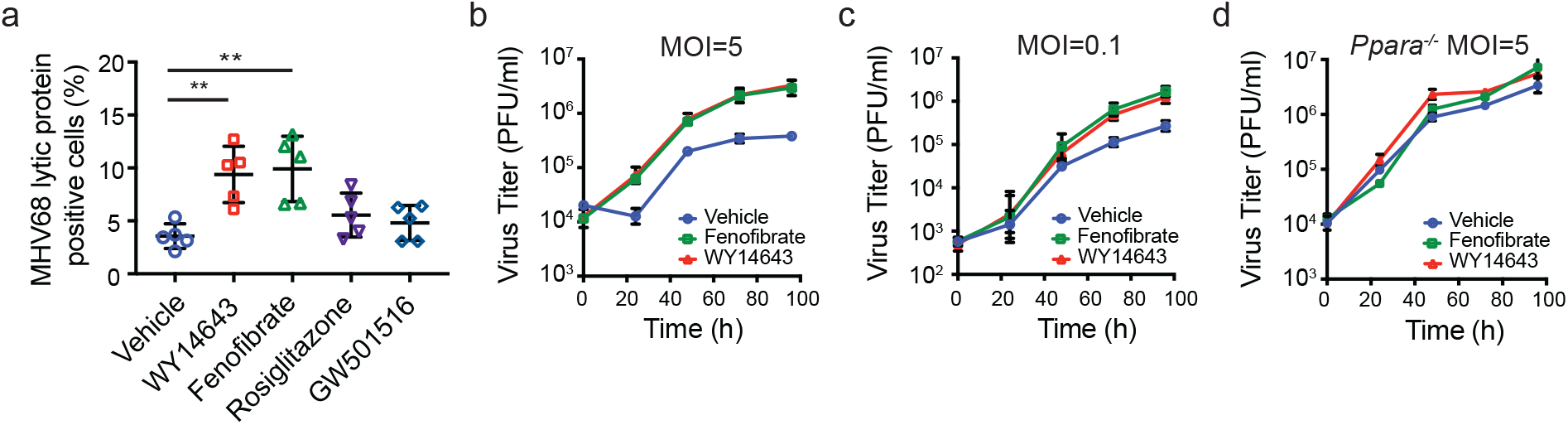
PPAR-α stimulation increases MHV68 replication. a, Macrophages from C57BL6/J mice were pretreated for 16 hours with WY14643, fenofibrate, rosiglitazone, or GW501516. Cells were infected with MHV68 at MOI=5. Quantification of cells expressing MHV68 lytic proteins using a polyclonal antibody 24 hours after infection was measured by flow cytometry. Data represents mean ± SD from 5 independent experimental. b-c, Growth curve of MHV68 in macrophages isolated from C57BL/6J mice after pretreatment with vehicle, WY14643 or fenofibrate. Cells were infected with MHV68 at MOI=5 (b) or MOI=0.1 (c). Virus quantitated by plaque assay on 3T12 cells. Data represents mean ± SD from 3 independent experiments. d, Growth curve of MHV68 in macrophages isolated from *Ppara^−/−^* mice after pretreatment with vehicle, WY14643 or fenofibrate. Cells were infected with MHV68 at MOI=5. Data represents mean ± SD from 3 independent experiments.

### PPAR-α stimulation suppresses type I IFN production in a STING-dependent manner

To broadly assess the impact of PPAR-α agonist on cells infected with virus, we performed RNA sequencing analysis on macrophages infected with virus and treated with WY14643. Six hours after infection, uninfected/vehicle treated, MHV68+/vehicle treated, uninfected/WY14643 treated cells, and MHV68+/WY14643 treated macrophages were prepared for RNA-seq. Pathway analysis of differentially expressed genes revealed that virus infection in both vehicle-treated and WY14643-treated cells increased expression of interferon responsive genes, genes involved in inflammation, and NFκB pathway genes (Figure 2a, Supplemental Table 1). However, PPAR-α stimulation reduced the magnitude of upregulation in infected cells. Thus, PPAR-α stimulation attenuates the early antiviral response in infected cells.

**Figure 2.**
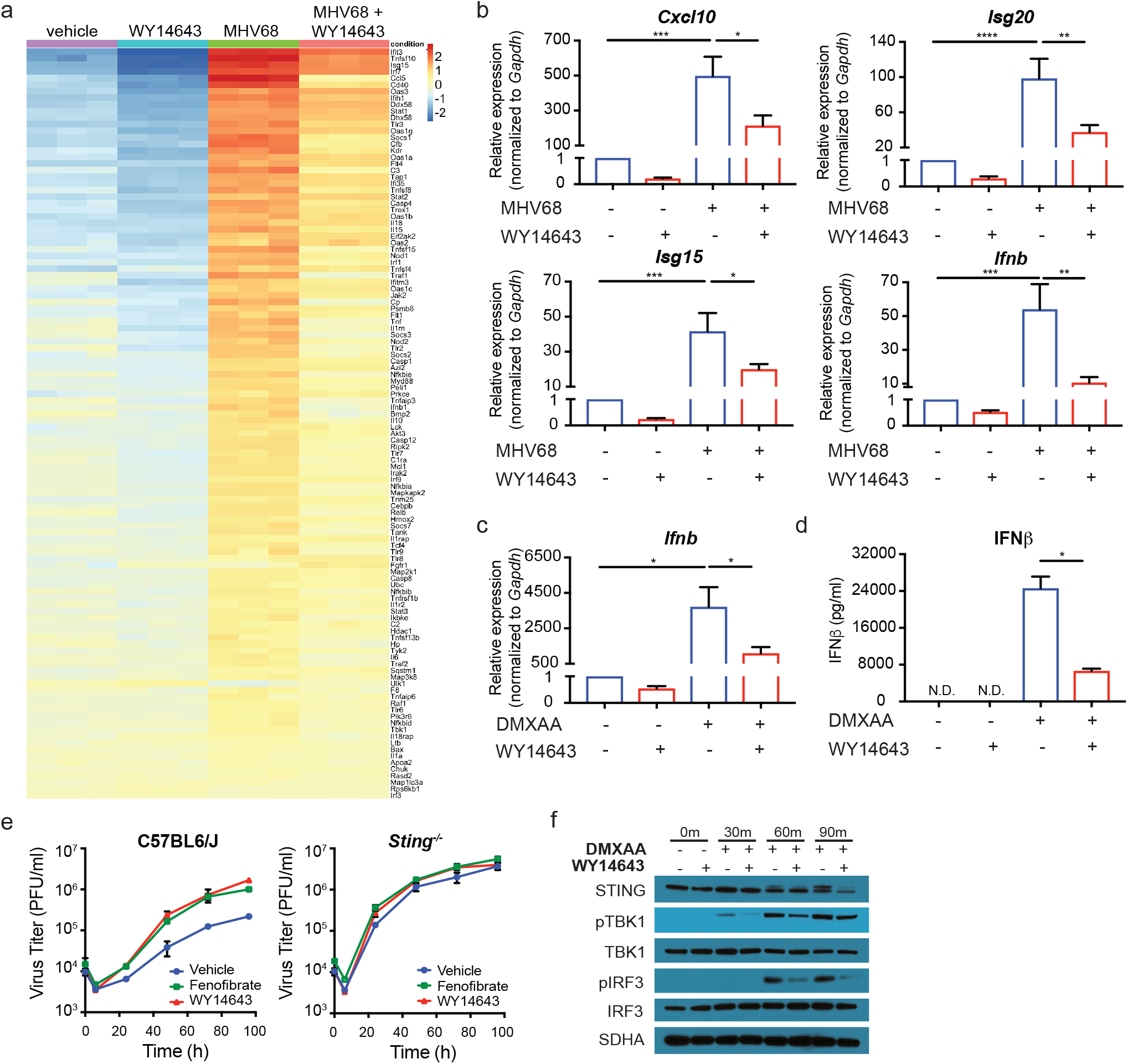
PPAR-α stimulation suppresses type I IFN in a STING-dependent manner. a, Macrophages were pretreated with vehicle or WY14643 for 16 hours prior to infection with MHV68, MOI=5. 6 hours after infection, RNA was isolated and prepared for RNA sequencing. b, Macrophages were pretreated with vehicle or WY14643 for 16 hours prior to infection with MHV68. qRT-PCR of *Cxcl10, Isg20, Isg15* and *Ifnb* before and 6 hours after MHV68 infection. Relative expression normalized to *Gapdh.* n=5 for *cxcl10,* n=7 for *Isg20,* n=6 for *Isg15,* n=8 for *Ifnb.* c, Macrophages were pretreated with vehicle or WY14643 for 16 hours prior to transfection with DMXAA. qRT-PCR of *Ifnb* before and 2 hours after transfection with DMXAA. Relative expression normalized to *Gapdh.* n=9. d, Macrophages were pretreated with vehicle or WY14643 for 16 hours prior to transfection with DMXAA. Concentration of IFNβ in culture medium before and 24 hours after transfection was determined by ELISA assay. n=3. e, Macrophages from C57BL6/J or *Sting^−/−^* mice were pretreated with vehicle or WY14643 for 16 hours prior to infection with MHV68, MOI=5. Virus growth was quantitated by plaque assay. n=3. f, Macrophages were pretreated with vehicle or WY14643 for 16 hours. Representative western blot of proteins involved in STING signaling before transfection and 30 mins, 60 mins, 90 mins after transfection with DMXAA. Data all shown as mean ± SD; *p<0.05, **p<0.01, ***P<0.001, statistical analysis was conducted using one-way ANOVA test.

To confirm the sequencing results, we quantified expression of a subset of interferon stimulated genes. We measured *Isg20, Isg15,* and *Cxcl10* and confirmed that expression of these genes is reduced in both uninfected and infected macrophages treated with WY14643 (Figure 2b). We also measured expression of *Ifnb* and found that expression is reduced after agonist treatment (Figure 2b). These data suggest that PPAR-α stimulation suppresses the interferon response at baseline and after viral infection.

Because of the suppression of the early antiviral response, we tested whether the effects of PPAR-α stimulation depended on the type I interferon response. To test this, we examined viral growth in wildtype and *Ifnar^−/−^* macrophages with and without agonist treatment. As expected, virus grew substantially more in *Ifnar^−/−^* cells compared with wildtype cells. PPAR-α stimulation did not further increase virus replication in the knockout cells (Supplemental Figure 3). Thus, PPAR-α stimulation effects depend on type I interferon.

MHV68 is a DNA virus that induces interferon downstream of the cGAS/STING pathway; therefore, we hypothesized that PPAR-α stimulation could antagonize the early induction of interferon after recognition of cytoplasmic DNA. Cytoplasmic DNA is sensed by cGAS, leading to activation of endoplasmic reticulum adapter molecule STING. STING phosphorylates TBK1 and IRF3, leading to transcription of IFNβ. IFNβ then signals through the type I IFN receptor to induce interferon stimulated gene expression and more IFNβ expression. To test if PPAR-α stimulation could suppress IFNβ induction after direct activation of STING, macrophages were treated with vehicle or agonist and stimulated with DMXAA, a murine agonist of STING. We found that PPAR-α stimulation suppressed IFNβ expression after DMXAA transfection (Figure 2c). As expected, the PPAR-γ and PPAR-δ agonists rosiglitazone and GW501516 did not suppress IFNβ expression after DMXAA treatment (Supplemental Figure 4). Moreover, PPAR-α stimulation suppressed production of IFNβ protein (Figure 2d). These data suggest that PPAR-α stimulation antagonizes the induction of interferon production downstream of the STING DNA sensing pathway.

To test if suppression of IFNβ depended on STING, we treated *Sting^−/−^* macrophages with PPAR-α agonist and infected with MHV68. We found that PPAR-α stimulation no longer increased virus replication in cells deficient in STING (Figure 2e). We examined activation of TBK1 and IRF3 and found that PPAR-α stimulation suppressed DMXAA-induced phosphorylation of both proteins (Figure 2f). These data indicate that the effects of PPAR-α agonist depend on STING expression and suggest that PPAR-α stimulation antagonizes the initial induction of IFNβ after recognition of cytoplasmic DNA.

### PPAR-α stimulation induces oxidative stress

We wondered if PPAR-α agonist effects could promote virus replication even if cells were treated with agonist after infection, or if the effects of agonist required pretreatment. To test this, we compared three different treatment protocols. We pretreated, as above, with PPAR-α agonist overnight and replaced agonist in the media following infection with MHV68. We compared this with pretreatment only (pre) or post treatment only (post). We found that pretreatment with PPAR-α agonist was required to increase MHV68 replication (Figure 3a). We found no increase in virus replication when cells were treated post-infection with agonist. We also found that pretreatment alone was sufficient to increase virus replication. These data suggest that PPAR-α stimulation is altering the cellular environment prior to infection in such a way that enhances virus replication.

**Figure 3.**
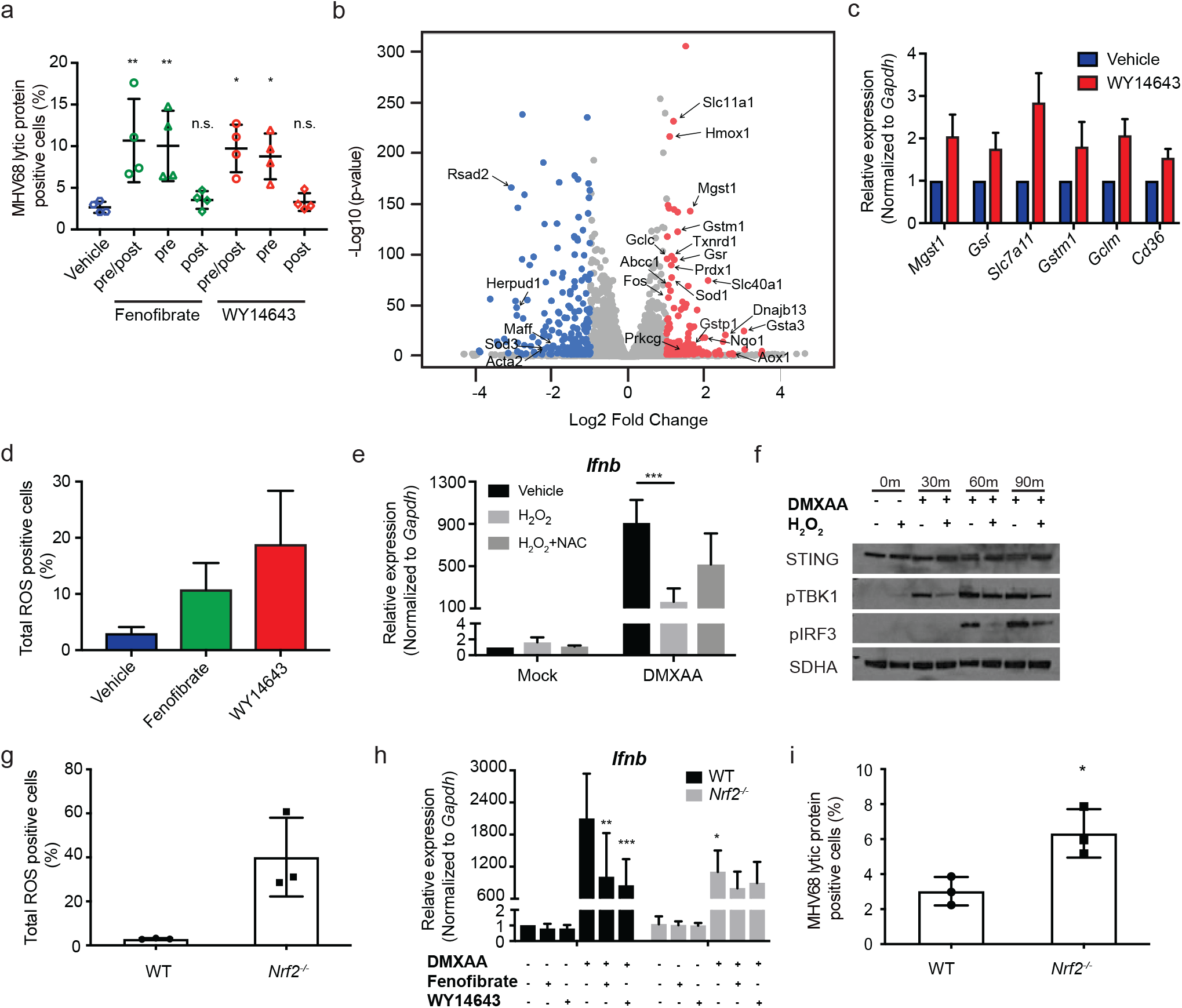
Oxidative stress inhibits STING activation and interferon production. a, Macrophages were treated with vehicle, WY14643 or fenofibrate agonist pre and post-infection (pre/post), pre-infection only (pre) or post-infection only (post). Quantification of cells expressing MHV68 lytic proteins by FACs assay 24 hours after infection. n=4. b, Comparison of vehicle-treated or WY14643-treated macrophages from RNA sequencing analysis in Figure 2A. c, Macrophages were treated with either vehicle control or WY14643 overnight. Genes associated with oxidative stress response were quantitated by qRT-PCR. Relative expression normalized to *Gapdh.* n=5 from independent experiments. d, Macrophages were treated with vehicle, fenofibrate or WY14643 for 6 hours. Quantification of total ROS in BMDMs by flow cytometry. n=7 with vehicle and fenofibrate treatment, n=6 with WY14643 treatment. e, Macrophages were pretreated with vehicle control or H_2_O_2_ for 20 mins. Half of the H_2_O_2_ samples were treated with NAC for 30 minutes and half were mock treated. Treatments were replaced with fresh media and cells were transfected with DMXAA. After 2 hours *Ifnb* was quantitated from lysates by qRT-PCR. n=4 from independent experiments. f, Macrophages were treated with either vehicle or H_2_O_2_ for 20 mins, and then transfected with DMXAA. Expression of STING, phosphorylated TBK1 (pTBK1), phosphorylated IRF3 (pIRF3), and SDHA was quantitated 0, 30, 60 or 90 minutes after transfection. Representative of 3 separate experiments. g, Macrophages were isolated from C57BL6/J or *Nrf2^−/−^* mice. Total ROS was quantitated by flow cytometry. n=3 from independent experiments. h, Macrophages isolated from WT or *Nrf2^−/−^* mice were pretreated for 16 hours with fenofibrate, WY14643, or vehicle. Cells were transfected with DMXAA and expression of *Ifnb* transcript was quantitated 2 hours later. Relative expression of *Ifnb* was normalized to *Gapdh.* n=5 Data all shown as mean ± SD; *p<0.05, **p<0.01, ***p<0.001, statistical analysis was conducted using one-way ANOVA or two-way ANOVA test. i, Macrophages isolated from WT or *Nrf2^−/−^* mice were infected with MHV68 at a MOI=5 for 24 hours. Cells expressing MHV68 lytic proteins were quantified by FACs assay. N=3 from independent experiments. Data all shown as mean ± SD; *p<0.05, **p<0.01, statistical analysis was conducted using oneway ANOVA or unpaired t-test.

In order to determine how PPAR-α stimulation was altering the cellular environment to promote virus replication and impair interferon induction, we examined vehicle and WY14643-only treatments from our sequencing data. PPAR-α agonist treated cells displayed increased antioxidant response pathways, indicating a possible increase in oxidative stress in PPAR-α agonist treated cells (Figure 3b, Supplemental Table 2). To confirm these results, we analyzed expression of antioxidant genes in untreated or PPAR-α agonist treated macrophages and confirmed increased expression of *Mgst1, Gsr, Slc7a11, Gstm1, Gclm,* and *Cd36* in agonist treated cells (Figure 3c). PPAR-α agonist treatment of macrophages also promoted increase total ROS (Figure 3d). These results are comparable to previous data indicating that PPAR-α agonist treatment increases β-oxidation of fatty acids in peroxisomes and enhances production of ROS^27,28^.

Given that PPAR-α stimulation increased antioxidant gene expression and ROS production and that these effects correlated with increased viral replication, we next tested whether increased ROS suppressed IFNβ production. Macrophages were treated with hydrogen peroxide (H_2_O_2_) and stimulated with DMXAA. H_2_O_2_ treatment suppressed DMXAA-induced IFNβ production (Figure 3e). In addition, N-Acetyl-L-cysteine (NAC), an antioxidant, partially neutralized the effects of H_2_O_2_ (Figure 3e). H_2_O_2_ treatment suppressed phosphorylation of TBK1 and IRF3 (Figure 3f). This indicates that increased oxidative stress suppresses the cGAS/STING pathway after activation with cytoplasmic DNA.

To determine if cells that have constitutive high levels of ROS have impaired interferon responses, we examined that antiviral response in nuclear factor E2-related factor 2 (NRF2) deficient cells. NRF2 is a transcription factor that regulates antioxidant responses by binding to antioxidant response elements (ARE) found in promoters of detoxication enzymes. NRF2 activity is regulated by KEAP1, and KEAP1 under non-stressed conditions promotes NRF2 ubiquitylation. When cells are under oxidative stress, NRF2 is released from KEAP1 and drives antioxidant gene expression^29^. *Nrf2^−/−^* cells do not induce antioxidant genes and have high levels of ROS^30–32^. We confirmed that *Nrf2^−/−^* macrophages had increased ROS (Figure 3g). When we stimulated *Nrf2^−/−^* macrophages with DMXAA they induced less *Ifnb* expression compared to wildtype cells (Figure 3H). Additionally, PPAR-α agonists did not further suppress *Ifnb* transcript production from knockout cells (Figure 3h). Together, these data suggest that ROS can suppress IFNβ downstream of direct STING activation.

To determine if the lower levels of IFNβ expressed in *Nrf2^−/−^* macrophages led to increased viral replication, we infected *Nrf2^−/−^* and wildtype macrophages with MHV68. We observed increase viral protein expression after infection in *Nrf2^−/−^* macrophages, indicating that virus replicated better in cells with higher levels of ROS and decreased IFNβ (Figure 3i).

### PPAR-*α* stimulation increases virus replication and lethality in mice

Because we found that PPAR-α stimulation *in vitro* suppressed interferon production, we hypothesized that agonist treatment of mice infected with MHV68 would alter herpesvirus replication. Previously published work established that mice deficient in the IFNa/β receptor (IFNAR) have increased viral replication and enhanced susceptibility to MHV68^33^. To determine if PPAR-α stimulation increased MHV68 replication *in vivo,* we injected wildtype and *Ppara^−/−^* mice with WY14643 or vehicle control for 7 days, starting 3 days prior to infection and continuing for 4 days after infection (Figure 4a). Mice were infected with a dose of MHV68 that does not cause lethality in wildtype mice treated with the drug vehicle. However, wildtype mice injected with PPAR-α agonist succumbed to infection (Figure 4b), at a frequency similar to type I interferon receptor deficient mice^33^. Moreover, *Ppara^−/−^* mice treated with agonist or vehicle all survived infection with MHV68, indicating that the lethality observed in wildtype mice treated with agonist is PPAR-α dependent (Figure 4c). Using a luciferase-tagged MHV68 (MHV68-M3FL) we imaged mice infected with virus over multiple days during acute infection^25,34^. We found that mice treated with WY14643 had increased virus replication compared to vehicle treated mice, and that this increase in virus replication was PPAR-α dependent (Figure 4d, e). These data suggest that PPAR-α stimulation significantly increased MHV68 acute replication.

**Figure 4.**
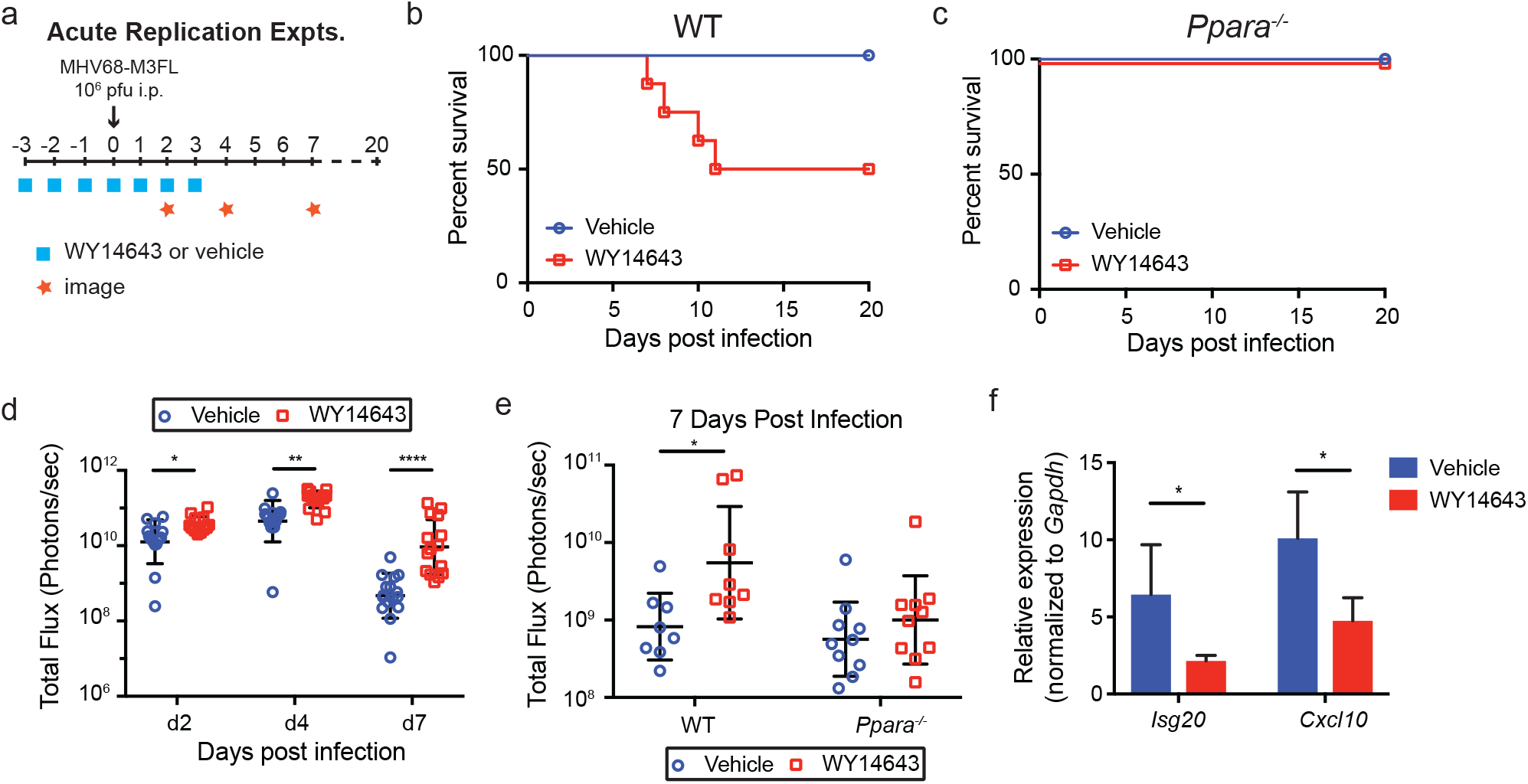
PPAR-α stimulation in mice increases virus replication 361 and lethality. a, Schematic for acute replication of MHV68 in mice. Mice were treated with either vehicle control (15% HS15) or WY14643 (100 mg/kg) for 7 days, starting 3 days before infection. Mice were infected intraperitoneally with MHV68-M3FL at dose of 10^6^ PFU. Acute replication of virus was measured at d2, d4 and d7 after infection using an IVIS bioluminescent imager. Survival of mice was monitored until 20 days after infection. b-c, Survival of mice infected with MHV68. C57BL6/J (b) or *Ppara^−/−^* (c) mice were treated with either vehicle control or WY14643, and infected with MHV68. n=8 in each group of C57BL6/J mice, n=10 in each group of *Ppara^−/−^* mice. Data shown as the pool of 2 independent experiments. d, Acute replication of MHV68 in mice measured by luminescent intensity. n=16 in vehicle treatment group, n=15 in WY14643 treatment group, data shown as the pool of 3 independent experiments. e, Acute replication of MHV68 in WT or *Ppara^−/−^* mice measured by luminescent intensity. n=8 in each group of WT mice, n=10 in each group of *Ppara^−/−^* mice. Data shown as the pool of 2 independent experiments. f, Peritoneal exudate cells were collected from vehicle or WY14643-treated mice 2 days after infection with MHV68. Relative expression of *Isg20* and *Cxcl10* was quantitated and normalized to *Gapdh.* n=4 from 2 independent experiments. Data all shown as mean ± SD; *p<0.05, **p<0.01, ***p<0.001, statistical analysis was conducted using one-way ANOVA or two-way repeated measures ANOVA test.

To determine if the interferon response was altered by PPAR-α stimulation, we analyzed interferon stimulated gene expression in the peritoneal cells for vehicle-treated and agonist-treated mice. We found that expression of *Isg20* and *Cxcl10* was decreased in agonist treated mice (Figure 4f), indicated that treatment with PPAR-α agonist during acute infection suppressed the interferon response.

## Discussion

We determined that activation of PPAR-α with clinically relevant agonists increases herpesvirus replication. In macrophages, PPAR-α stimulation increased virus replication and suppressed the induction of an interferon response in a STING-dependent manner. PPAR-α stimulation induced oxidative stress in cells and high levels of oxidative stress impaired STING activation. This is the first evidence that PPAR-α activation and peroxisomal metabolism regulate the cytoplasmic DNA-sensing pathway and IFNβ production downstream of STING. PPAR-α stimulation significantly increased herpesvirus replication in mice, leading to increased lethality of agonist-treated mice.

Metabolism is recognized as an important regulator of immunity. Metabolic reprogramming of macrophages, in particular, is essential for modifying their inflammatory phenotype and function. Glycolysis drives inflammatory macrophage responses, whereas high fatty acid oxidation was long thought to support anti-inflammatory macrophage responses^35^. It is now clear that fatty acid oxidation is not just anti-inflammatory, because oxidation of palmitate promotes the production of mitochondrial ROS, which induces inflammasome activation^36,37^. Moreover, mitochondrial ROS promotes the production of the proinflammatory cytokines IL-1β, IL-6, and TNFa following LPS stimulation of macrophages^38^. Our data adds another level of complexity to the role of ROS in inflammation. Our data indicates that ROS, perhaps derived from peroxisomal β-oxidation of very long chain fatty acids or other sources, impairs STING activation and interferon production. Further work is needed to determine the source of ROS. However, our data thus far suggests that the induction of peroxisomal metabolism and fatty acid oxidation impairs innate immune signaling.

These data reveal that peroxisomal metabolism and increased ROS production regulates the cytoplasmic DNA sensing pathway. The adapter molecule MAVS that signals downstream of RNA recognition is localized to mitochondria and peroxisomes. Interestingly, the peroxisomal localization of MAVS dictates unique signaling pathways and gene responses distinct from mitochondrial MAVS^39,40^. Together these data along with our data indicate that peroxisomes are critical organelles for the induction of innate immune signals, either through localization of signaling molecules or production of metabolites that regulate immune signaling.

Herpesviruses use multiple mechanisms to maintain chronic infections in the host and peroxisomal metabolism is a putative mechanism. To avoid clearance, herpesviruses evade immune recognition by downregulating antigen presentation, expressing viral proteins that regulate the interferon pathway, inhibiting cell cycle arrest and apoptosis, and encoding various molecular mimics. This is indicative of a long evolutionary relationship with the host. It also suggests that herpesviruses modify numerous pathways within the host cell to tune the balance between latency and reactivation. Peroxisomal proliferation is a feature of herpesvirus infection, but the function of peroxisomal biogenesis during latency has been elusive. A recent report found that HCMV and HSV-1 induced peroxisome biogenesis and plasmalogen production, leading to enhanced virus envelopment and lytic replication^20^. However, these data do not address the potential role of peroxisome biogenesis in chronic infection with a herpesvirus. KSHV latent cell survival requires peroxisomal lipid metabolism, suggesting that KSHV regulates peroxisomal lipid metabolism to support latent infection^19^. One hypothesis is that peroxisomes and the redox balance of the cell control latency by adjusting the production of type I interferon.

A key remaining question is how ROS regulates STING activation. Recent structural data provide numerous insights into the mechanism of cGAMP activation of STING^41,42^. STING dimers in the ER form polymers upon binding to cGAMP and disulfide bonds in the cytosolic domain stabilize STING polymers. These inter-dimer crosslinks are important for STING activation. One possibility is that increased oxidative stress leads to oxidation of cysteine residues in the cytosolic domain, which interferes with the polymerization and activation of STING^43^.

Overall, our work suggests that activation of PPAR-α suppresses interferon and antiviral immunity. Moreover, these data indicate that metabolism and particularly ROS have both proinflammatory and anti-inflammatory functions. These results have implications for therapies that target PPAR-α or oxidative stress pathways for the treatment of infection, metabolic disorders, cancer and autoimmune disease.

## Acknowledgments

We thank members of the Reese and Yan labs for technical assistance, David Mangelsdorf and Steven Kleiwer for reagents and expertise, Dr. Wenhan Zhu for help with data analysis, and Drs. Julie Pfeiffer and Lora Hooper for editorial assistance. We also thank the UTSW Flow Cytometry core and the Genomic and Microarray core for technical assistance. The Reese lab is supported by the American Heart Association (17SDG33670071), the NIH (1R01AI130020-01A1), and the Pew Scholars Program.

## Author Contributions

L.T. performed and analyzed experiments, with technical assistance from A.L, G.W and T.C. I.D. performed RNAseq analysis. N.Y. provided reagents and expertise. T.A.R. directed the project and wrote the manuscript.

---

**Supplemental Figure 1.**
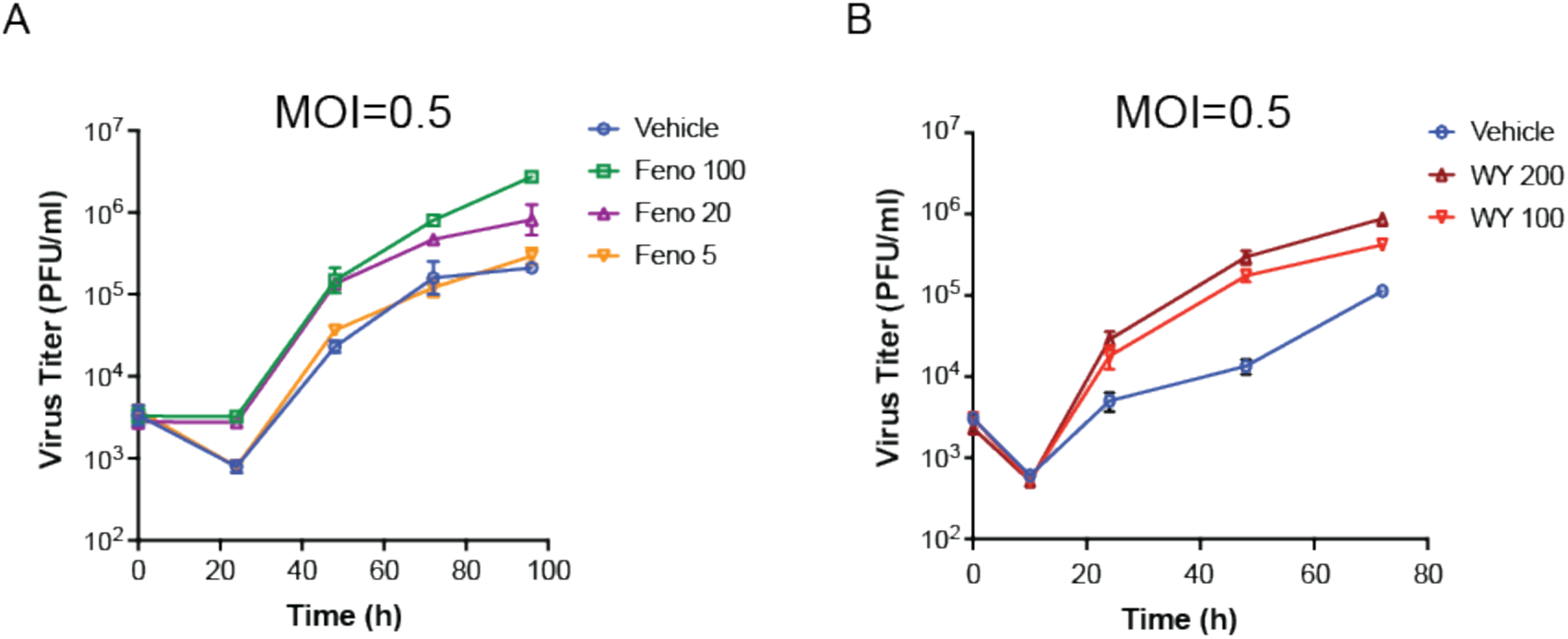
Viral replication with different doses of fenofibrate and WY14643. (A) Macrophages from C57BL6/J mice were treated with vehicle or increasing doses of fenofibrate (5, 20, or 100 μM) for 16 hours. Cells were infected with MHV68 at MOI=0.5 and viral growth was quantitated by plaque assay on 3T12 cells. (B) Macrophages from C57BL6/J mice were treated with vehicle or increasing doses of WY14643 (100 or 200 μM) for 16 hours. Cells were infected with MHV68 at MOI=0.5 and viral growth was quantitated by plaque assay on 3T12 cells.

**Supplemental Figure 2.**
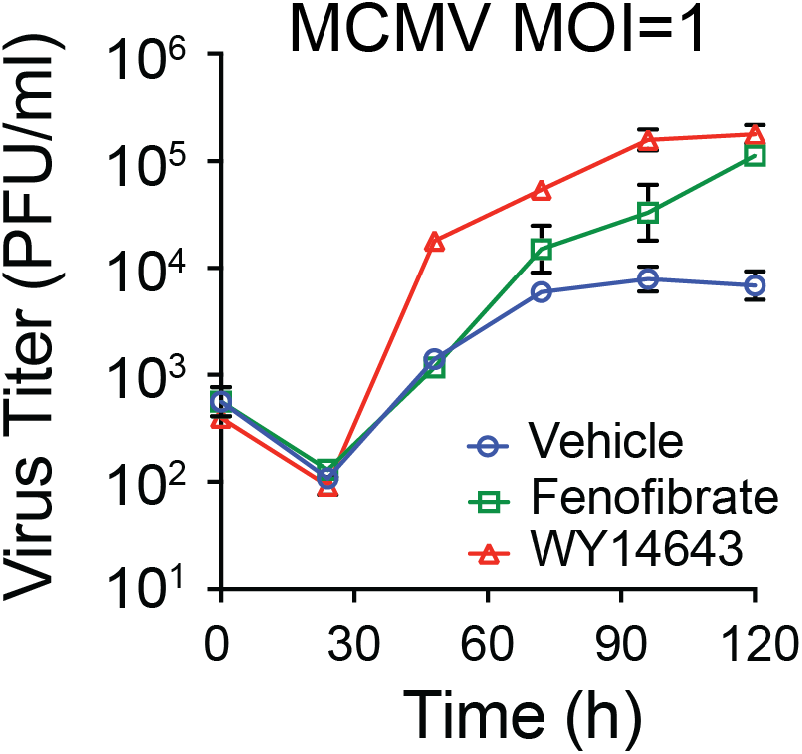
PPAR-α stimulation increases virus replication of MCMV. Growth curve of MCMV in macrophages isolated from C57BL/6J mice after pretreatment with vehicle, WY14643 or fenofibrate. Cells were infected with MCMV at MOI=1. Data represents mean ± SD from 2 independent experiments.

**Supplemental Figure 3.**
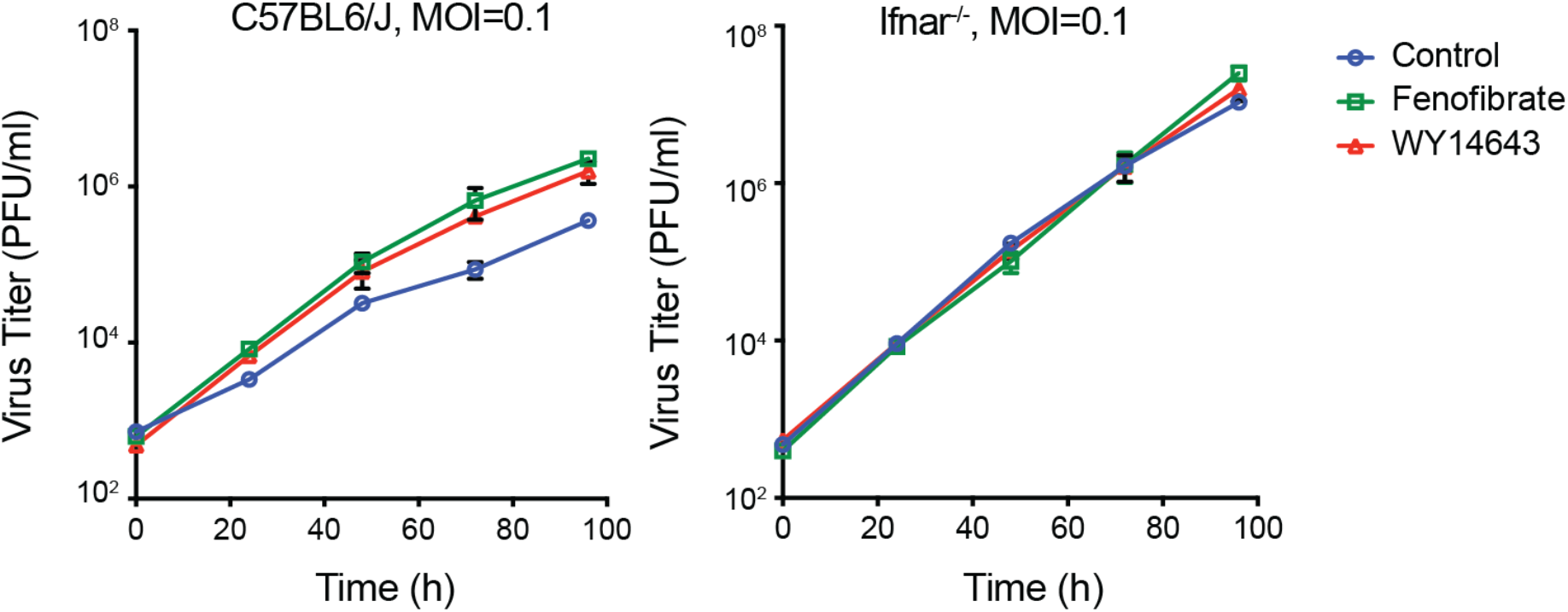
Effects of PPAR-α stimulation are IFNAR-dependent. Macrophages from C57BL6/J mice or *Ifnar^−/−^* mice were treated with vehicle, fenofibrate or WY14643 for 16 hours. Cells were infected with MHV68 at MOI=0.1 and viral growth was quantitated by plaque assay on 3T12 cells.

**Supplemental Figure 4.**
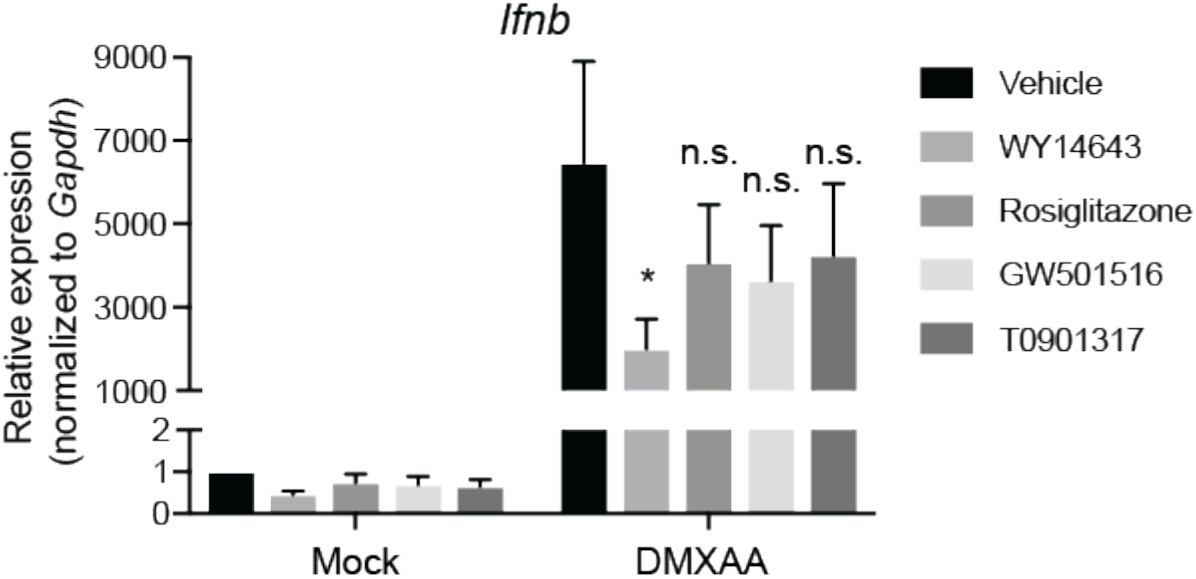
*Ifnb* expression after treatment with PPAR-α, PPAR-γ, PPAR-δ, or LXR-a agonists. Macrophages from C57BL6/J mice were treated with vehicle, WY14643, rosiglitazone, GW501516, or T0901319 for 16 hours. Cells were then transfected with DMXAA and *Ifnb* expression was quantitated from lysates 2 hours after transfection. Expression was normalized to *Gapdh.* Data all shown as mean ± SD; *p<0.05, n.s. (not significant), relative to DMXAA/vehicle treated. Statistical analysis was conducted using two-way ANOVA test.

## Material and Methods

### Animals

C57BL/6J, B6;129S4-*Pparα*^tm1Gonz^/J, C57BL/6J-*Tmem173*^gt^/J B6(Cg)-*Tmem173*^tm1.2Camb^/J, B6.129S2-/*Ifnar1*^tm1Agt^/Mmjax, and B6.129X1-*Nfe2/2*^tm1Ywk^/J were purchased from The Jackson Laboratory. All mice were housed under specific pathogen-free, double-barrier facility at the University of Texas Southwestern Medical Center. Mice were fed autoclaved rodent feed and water. Mice were maintained and used under a protocol approved by UT Southwestern Medical Center Institutional Animal Care and Use Committee (IACUC).

### Chemicals

For *in vitro* experiments, PPARα agonists, Fenofibrate and WY14643; PPARβ agonist, GW501516; PPARγ agonist, Rosiglitazone were purchased from Sigma. For *in vivo* experiments, WY14643 was purchased from Cayman. Hydrogen peroxide solution and N-Acetyl-L-Cysteine was purchased from Sigma.

### Cell culture

Bone marrow derived macrophages were differentiated in DMEM (Corning) with 10% FBS supplemented with 1% glutamine (Corning), 1% HEPES (Corning) and 10% CMG14 supernatant for 7 days^1^. 3T12 cells were maintained in DMEM with 5% FBS supplemented with 1% glutamine and 1% HEPES.

### Generation of virus stocks

Murine γ-herpesvirus 68 (WUSM stain) was purchased from ATCC. Murine γ-herpesvirus 68-M3FL was generated as previously reported^2^.

### Virus infection

Fully differentiated BMDMs were seeded on 24 well plates (1.5×10^5^ cells per well) or 6 well plate (10^6^ cells per well). Cells were pretreated with either vehicle control (0.1% DMSO) or agonists (Fenofibrate 50 μM, WY14643 200 μM, Rosiglitazone 1 μM, GW501516 100 nM) for 16 hours. The next days, macrophages were infected with MHV68 at multiplicity of infection (MOI)=5 or 0. 1. For MCMV experiments, cells were infected at MOI=1. After an hour, cells were washed with PBS twice to remove unabsorbed viruses and resuspended in medium containing treatments. For growth curve, samples were collected at 0 hour, 24 hours, 48 hours, 72 hours and 96 hours after infection and were frozen at −80 °C. The titer of virus was determined by plaque assay in 3T12 cells. For flow cytometric analysis, cells were collected 24 hours after infection.

### Flow cytometry for MHV68 lytic proteins positive cells

To determine the percentage of cells that expressing lytic proteins of MHV68 infection, cells were harvested 24 hours after infection, and fixed with 2% formaldehyde, blocked with 10% mouse serum and 1% Fc block (CD16/32, BioLegend), then stained with polyclonal rabbit antibody to MHV68 (1:1000)^3,4^, followed by secondary goat anti-rabbit Alexa Fluor-647 (Invitrogen).

### Transfection

Cells were seeded on 6 well plates and treated as needed. Cells were transfected with STING ligand (DMXAA, 10 μg/ml, InvivoGen) using Lipofectamine 3000 (Thermo Fisher Scientific) according to the manufacturer’s protocol.

### Plaque assay

The concentration of virus was titered in 3T12 cells. The frozen samples containing viruses were thawed in incubator. The samples were serial diluted, then added to a monolayer of 3T12 cells. After an hour of absorption, the cells were then covered with 1% methylcellulose. Plates were incubated at 37 °C for 7 days, and the plaques were stained with 0.1% crystal violet.

### Western blot

Cells were lysed with RIPA buffer (150 mM NaCl, 1% NP-40, 0.5% sodium deoxycholate, 0.1% SDS, 25 mM Tris with protease inhibitor cocktail). Protein concentration were determined using Bradford assay (Bio-Rad). Equal amounts of protein were mixed with 5×loading sample buffer, resolved by 4-12% Bis-Tris plus gels (Thermo Fisher Scientific), and transferred to nitrocellulose membrane. Proteins were labeled with primary antibodies against STING (1:1000, Catalogue no.13647S, Cell Signaling), TBK1 (1:1000, Catalogue no. 3504S, Cell Signaling), IRF3 (1:1000, Catalogue no. 4302S, Cell Signaling), pTBK1 (1:1000, Catalogue no. 5483S, Cell Signaling), pIRF3 (1:1000, Catalogue no. 4947S, Cell Signaling), PPARa (1:1000, Catalogue no. Sc-398394, Santa Cruz), SDHA (1:5000, Catalogue no. ab14715, Abcam), β-actin (1:5000, Catalogue no. A2228, Sigma). Secondary antibodies used are donkey-anti-rabbit (1:5000, Catalogue no.711-035-152, Jackson ImmunoResearch Laboratory) and goat-anti-mouse peroxidase (1:5000, Catalogue no.115-035-174 Jackson ImmunoResearch Laboratory). Membranes were developed using Luminata Forte Western HRP substrate (Millipore).

### RT-qPCR

BMDMs in 6 well plates were either infected with MHV68 at MOI=5 for 6 hours or transfected with STING ligand DMXAA at 10 μg/ml for 2 hours. RNA was extracted using Qiagen RNeasy Mini Kit (Qiagen) and reverse transcribed into cDNA using SuperScript VILO cDNA Synthesis Kit (Thermo Fisher Scientific). Relative quantification of target genes was determined using PowerUp SYBR Green Master Mix (Thermo Fisher Scientific) in a QuantStudio 7 Flex real time PCR system. Primers used for amplifying target genes are listed in Table 3.

**Table 1.**
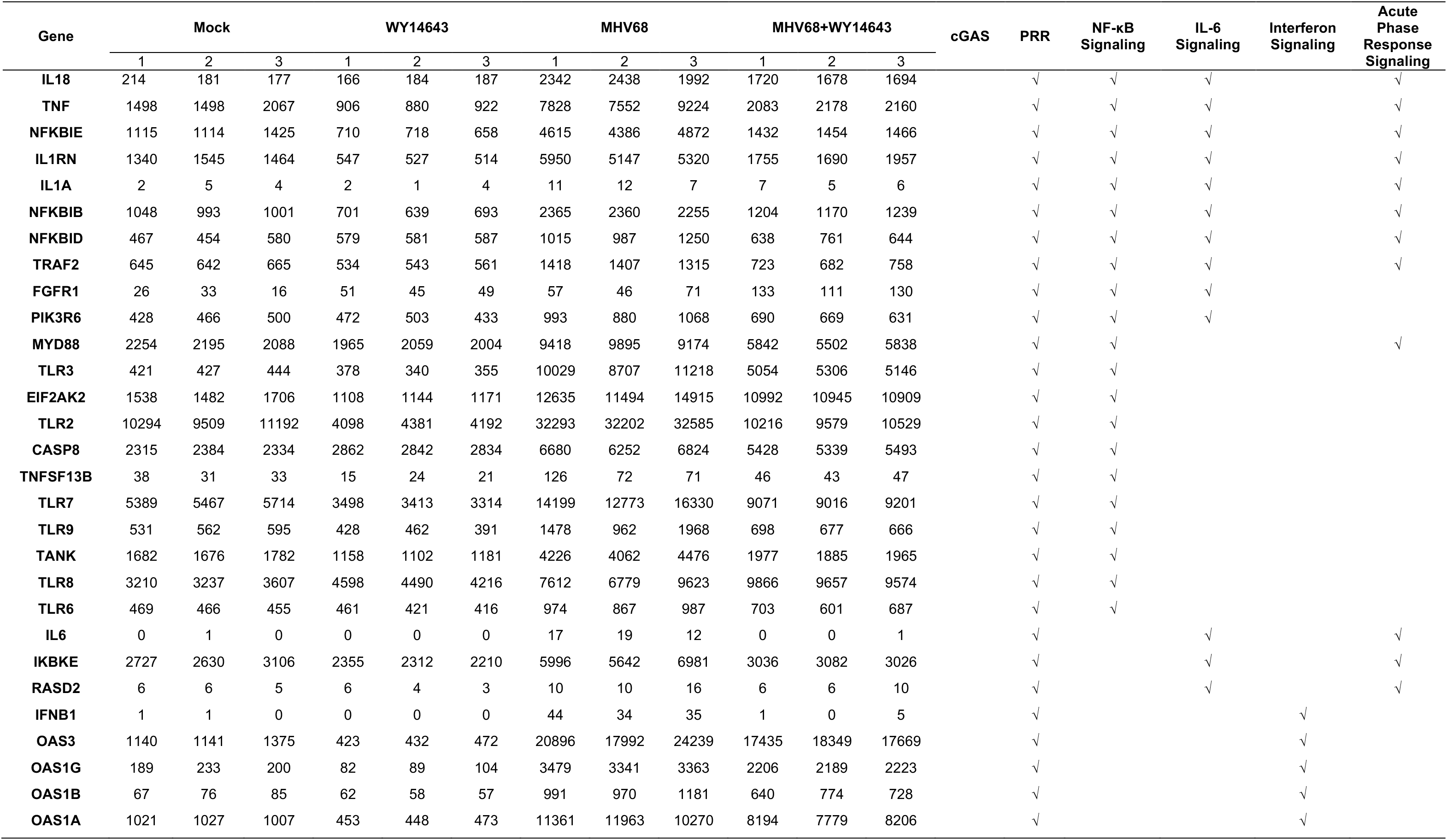
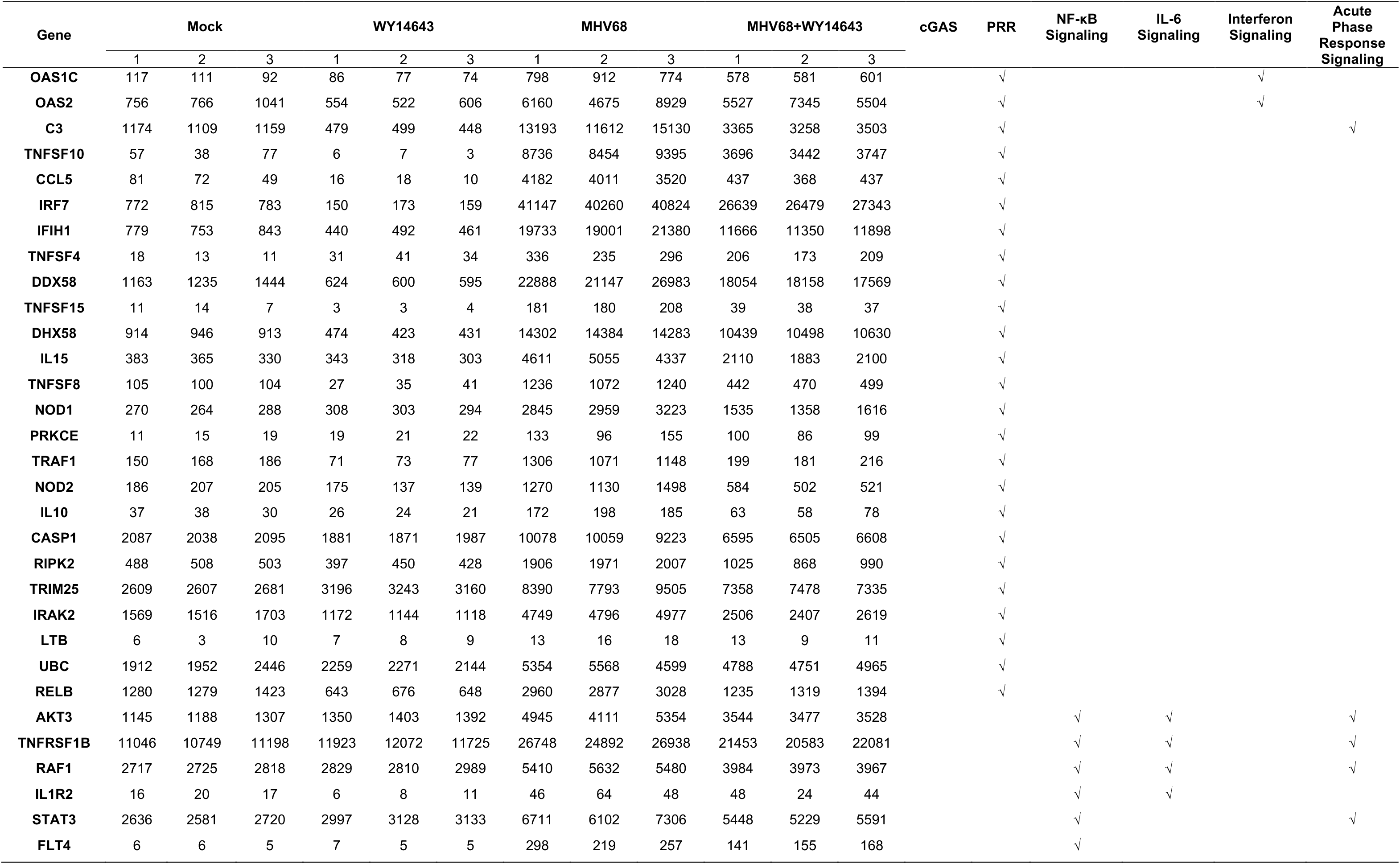
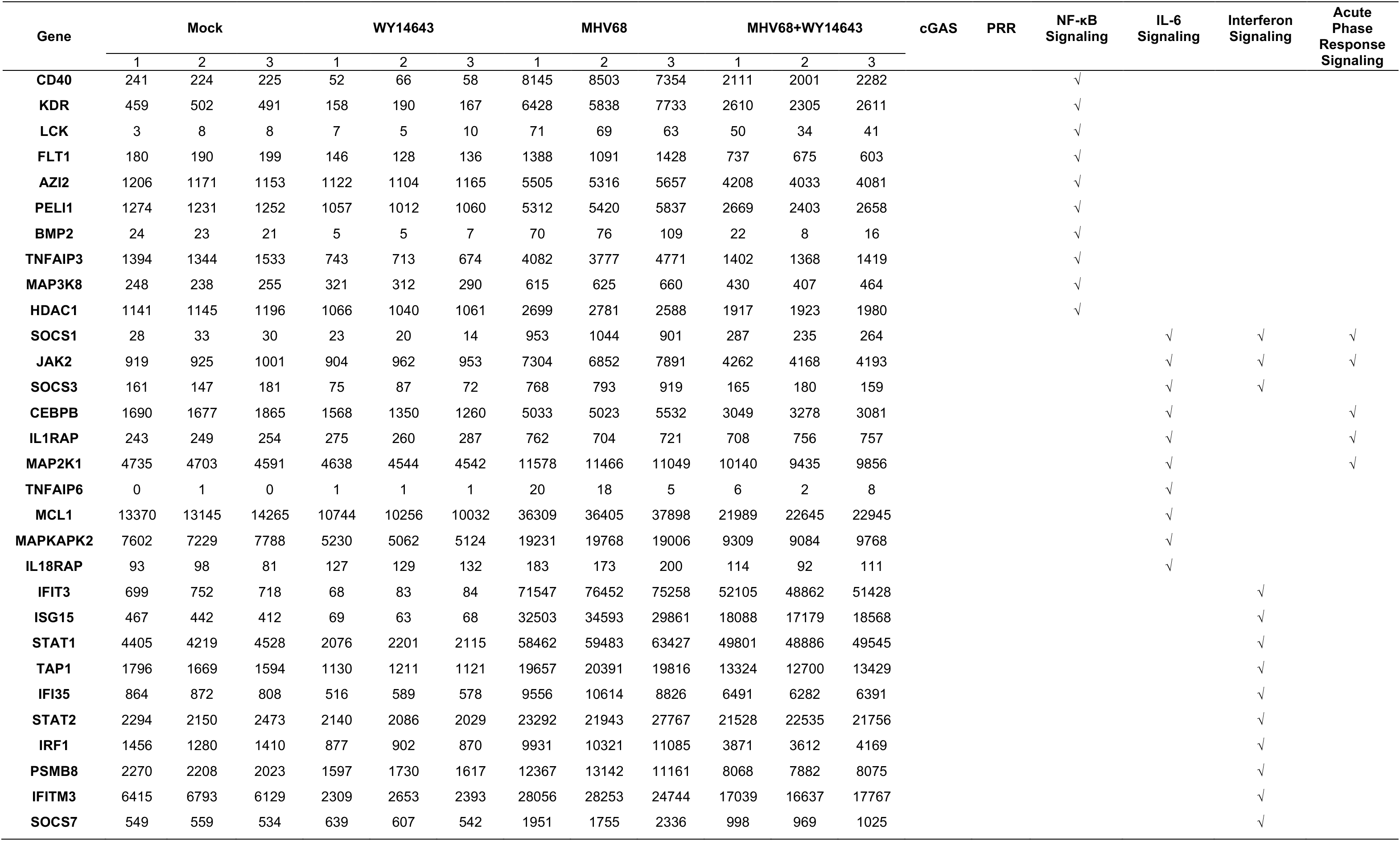
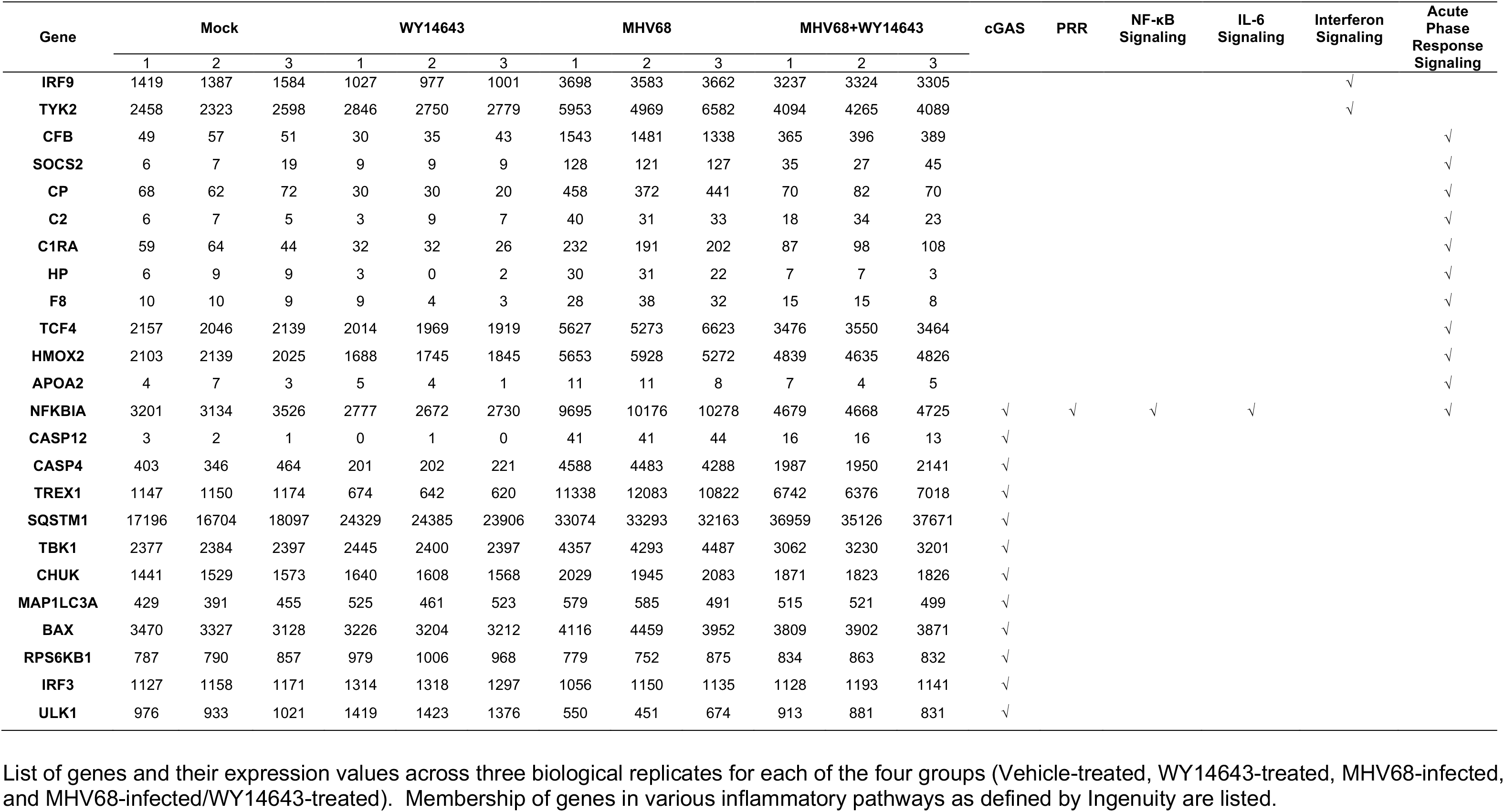
Differential expression of genes involved in innate inflammatory response

**Table 2.**
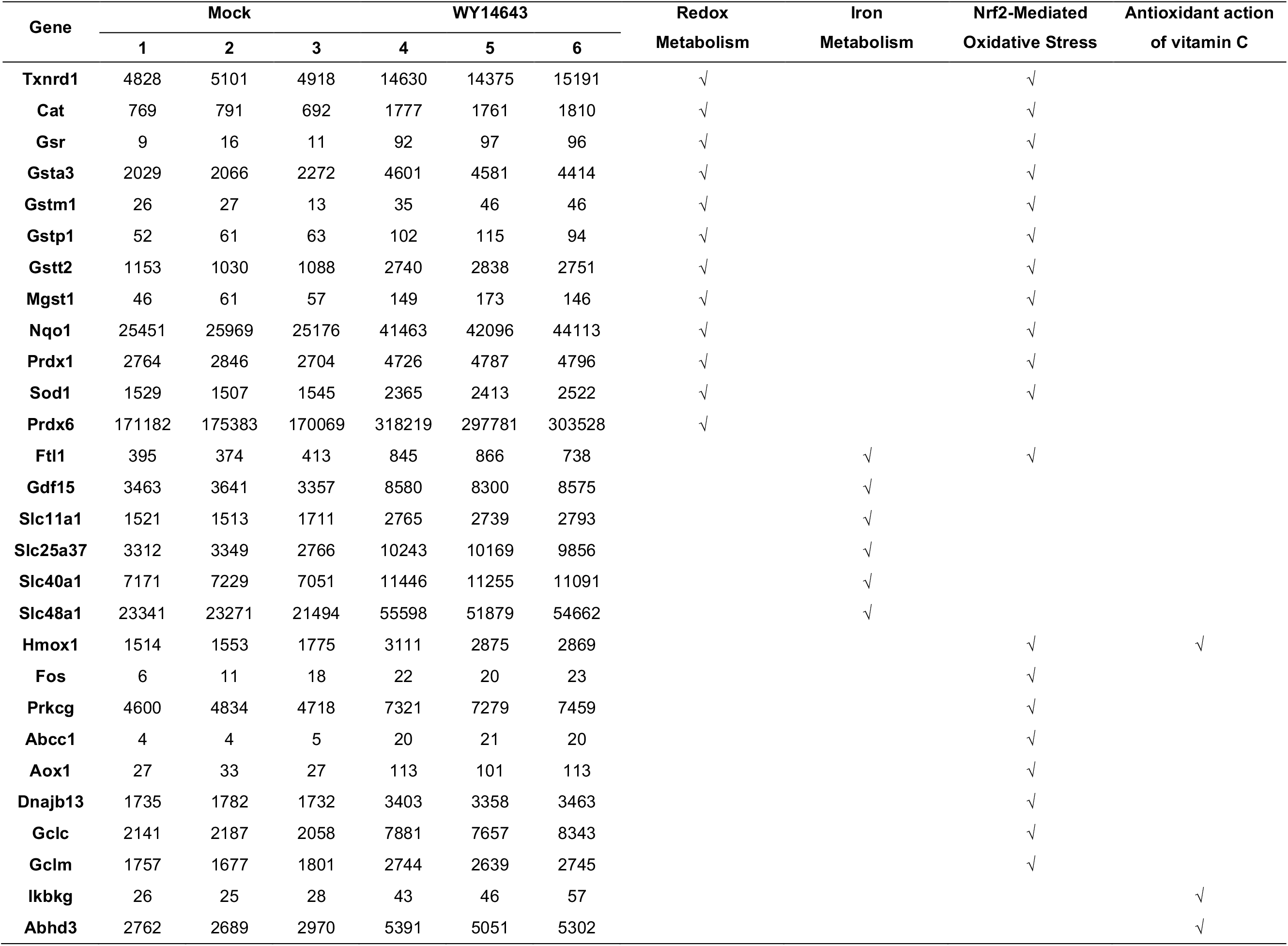
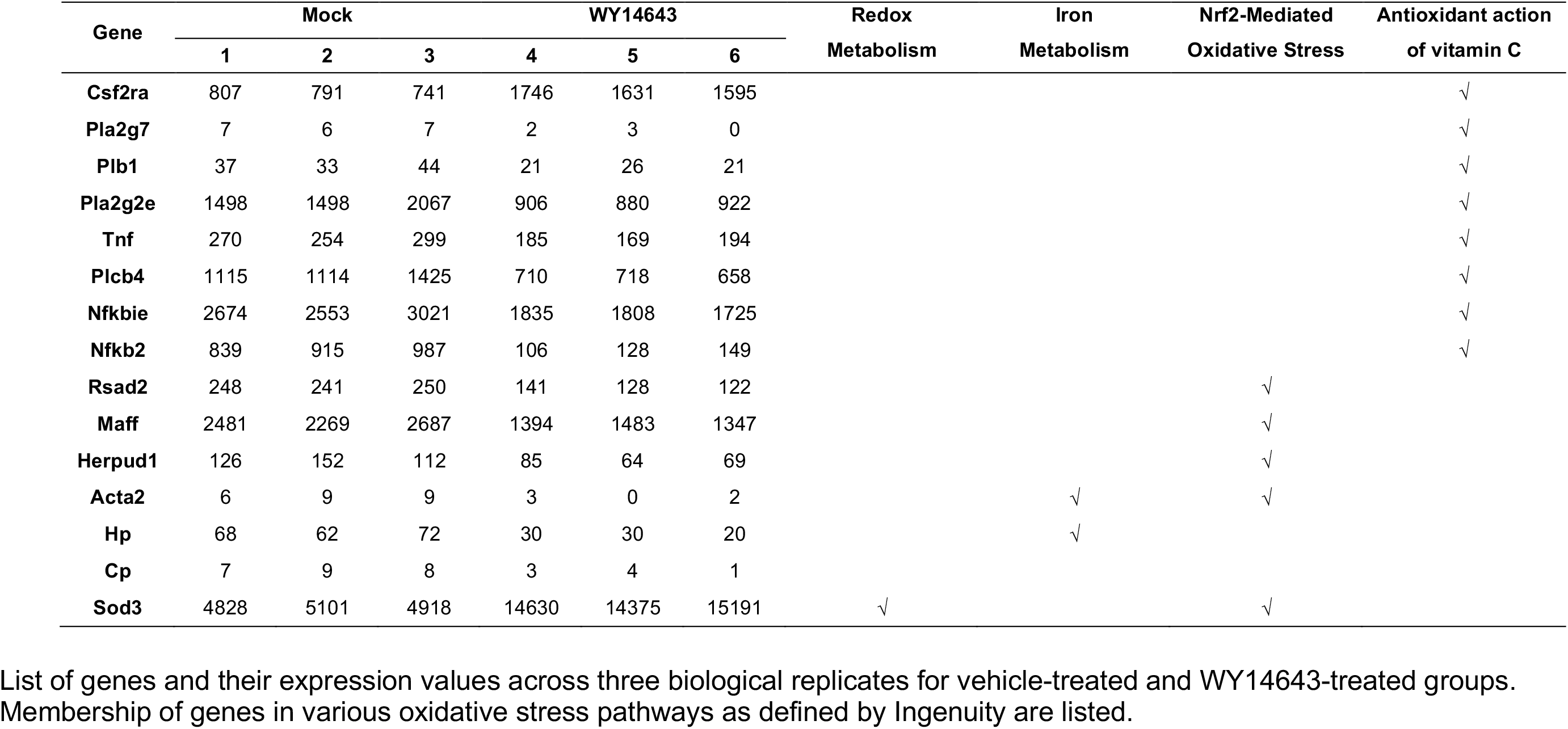
Differential expression of genes involved in cellular oxidative stress with WY14643 treatment

**Table 3.**
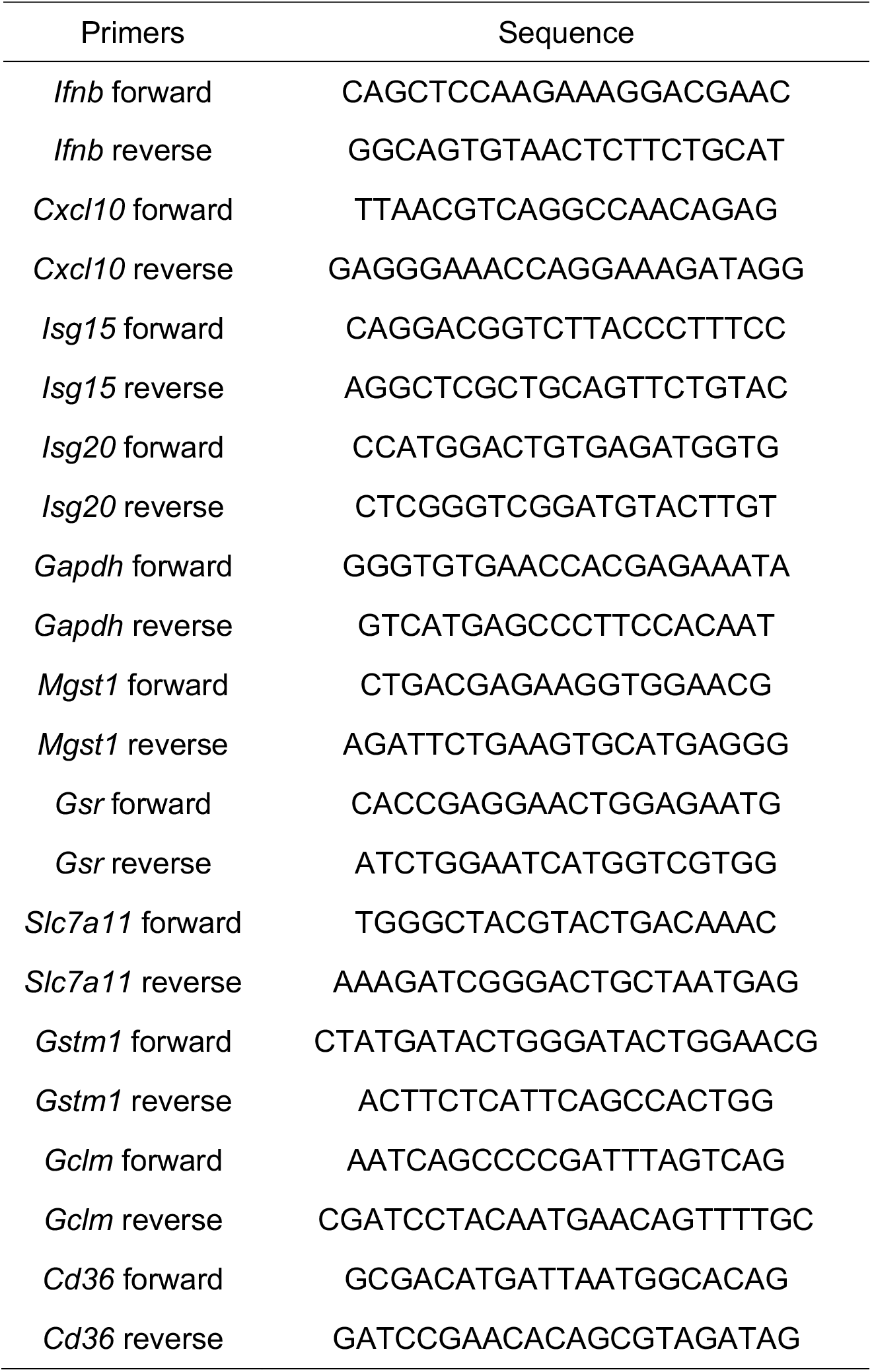
QPCR primers for target genes

### IFNβ ELISA Assay

BMDMs were seeded in 24 well plates. The next day, cells were treated with either vehicle control or WY14643 for 16 hours. Cells were then transfected with 10 μg/ml DMXAA. Supernatant of cells were collected 24 hours after transfection and frozen at −80 °C. The concentration of IFNβ in the supernatant was determined with PBL IFN Beta ELISA Kit (PBL Assay Science).

### Total ROS measurement

The total ROS production was determined using ROS-ID Total ROS detection kit (Enzo) according to the manufacture’s protocol. Cells were seeded on 24 well plates, and treatment with indicated compounds or DMSO for 6 hours, followed by incubation with oxidative stress detection reagent for 30 mins at 37 °C. The fluorescence of the detection reagent upon oxidation was analyzed by flow cytometry using a FACSCalibur (BD Biosciences).

### MHV68 acute replication in mouse

Experiments were carried out using 8-12 weeks old male mice under the protocol approved by IACUC. Mice were injected intraperitoneally with either vehicle control (15% HS15 in normal saline) or WY14643 (100 mg/kg) for 1 week starting from 3 days before virus infection. Mice were then infected with MHV68-M3FL at the dose of 10^6^ PFU through intraperitoneal route^2,4^. To quantify virus-encoded luciferase expression, mice were weighed and injected with 150 mg/kg of D-Luciferin (GOLDBIO) prior to imaging using IVIS Lumina III In Vivo Imaging System (PerkinElmer). Total flux (Photons/second) of the abdominal region was determined using Living Image software (PerkinElmer). Survival of the mice were recorded until 20 days after infection.

### RNA seq and data analysis

RNA samples were extracted with RNeasy Mini Kit (Qiagen) and RNA library was prepared using TruSeq Stranded mRNA Sample Preparation Kit (Illumina). Samples were sequenced on NextSeq 500/500 sequencer (Illumina) with SE-85. RNA seq data is normalized and analyzed based on the use of “internal standards”^5^ that characterize some aspects of the system’s behavior, such as technical variability, as presented elsewhere^6,7^. The two-step normalization procedure and the Associative analysis functions are implemented in MATLAB (MathWorks, MA) and available from authors upon request. Functional analysis of identified genes was performed with Ingenuity Pathway Analysis (IPA; Ingenuity System). The sequencing data has been deposited in the European Nucleotide Archive under the accession number PRJEB33753.

## QUANTIFICATION AND STATISTICAL ANALYSIS

All data are presented as mean ± SD. Statistical comparisons were performed using GraphPad Prism 7.0 software. Data were compared using unpaired two-tailed t test, one-way or two-way ANOVA. Statistical significance was set at p < 0.05. The numbers of independent replicates (n) are reported in the figure legends.

## References

1. Grygiel-Górniak, B. Peroxisome proliferator-activated receptors and their ligands: nutritional and clinical implications--a review. Nutrition journal 13, 17 (2014).

2. Staels, B. et al. Mechanism of action of fibrates on lipid and lipoprotein metabolism. Circulation 98, 2088–2093 (1998).

3. Staels, B. & Fruchart, J.-C. C. Therapeutic roles of peroxisome proliferator-activated receptor agonists. Diabetes 54, 2460–70 (2005).

4. Bensinger, S. J. & Tontonoz, P. Integration of metabolism and inflammation by lipid-activated nuclear receptors. Nature 454, 470–477 (2008).

5. Straus, D. S. & Glass, C. K. Anti-inflammatory actions of PPAR ligands: new insights on cellular and molecular mechanisms. Trends in immunology 28, 551–558 (2007).

6. Husson, M.-O. O. et al. Modulation of host defence against bacterial and viral infections by omega-3 polyunsaturated fatty acids. The Journal of infection 73, 523–535 (2016).

7. Jones, G. J. & Roper, R. L. The effects of diets enriched in omega-3 polyunsaturated fatty acids on systemic vaccinia virus infection. Scientific reports 7, 15999 (2017).

8. Manoharan, I. et al. Homeostatic PPARα Signaling Limits Inflammatory Responses to Commensal Microbiota in the Intestine. Journal of immunology (Baltimore, Md.: 1950) 196, 4739–4749 (2016).

9. Anderson, M. & Fritsche, K. L. (n-3) Fatty acids and infectious disease resistance. The Journal of nutrition 132, 3566–76 (2002).

10. Irons, R., Anderson, M. J., Zhang, M. & Fritsche, K. L. Dietary fish oil impairs primary host resistance against Listeria monocytogenes more than the immunological memory response. The Journal of nutrition 133, 1163–9 (2003).

11. O’Neill, L. A., Kishton, R. J. & Rathmell, J. A guide to immunometabolism for immunologists. Nature Reviews Immunology 16, 553–565 (2016).

12. Forrester, S. J., Kikuchi, D. S., Hernandes, M. S., Xu, Q. & Griendling, K. K. Reactive Oxygen Species in Metabolic and Inflammatory Signaling. Circulation research 122, 877–902 (2018).

13. Soucy-Faulkner, A. et al. Requirement of NOX2 and reactive oxygen species for efficient RIG-I-mediated antiviral response through regulation of MAVS expression. PLoS pathogens 6, e1000930 (2010).

14. Kong, X., Thimmulappa, R., Kombairaju, P. & Biswal, S. NADPH oxidase-dependent reactive oxygen species mediate amplified TLR4 signaling and sepsis-induced mortality in Nrf2-deficient mice. Journal of immunology (Baltimore, Md.: 1950) 185, 569–77 (2010).

15. Kobayashi, E. H. et al. Nrf2 suppresses macrophage inflammatory response by blocking proinflammatory cytokine transcription. Nature communications 7, 11624 (2016).

16. Agod, Z. et al. Regulation of type I interferon responses by mitochondria-derived reactive oxygen species in plasmacytoid dendritic cells. Redox biology 13, 633–645 (2017).

17. Yu, Y., Clippinger, A. J. & Alwine, J. C. Viral effects on metabolism: changes in glucose and glutamine utilization during human cytomegalovirus infection. Trends in microbiology 19, 360–367 (2011).

18. Vastag, L., Koyuncu, E., Grady, S. L., Shenk, T. E. & Rabinowitz, J. D. Divergent effects of human cytomegalovirus and herpes simplex virus-1 on cellular metabolism. PLoS pathogens 7, e1002124 (2011).

19. Sychev, Z. E. et al. Integrated systems biology analysis of KSHV latent infection reveals viral induction and reliance on peroxisome mediated lipid metabolism. PLoS pathogens 13, e1006256 (2017).

20. Beltran, P. M. et al. Infection-Induced Peroxisome Biogenesis Is a Metabolic Strategy for Herpesvirus Replication. Cell host & microbe 24, 526–541.e7 (2018).

21. Choi, Y. B., Choi, Y. & Harhaj, E. W. Peroxisomes support human herpesvirus 8 latency by stabilizing the viral oncogenic protein vFLIP via the MAVS-TRAF complex. PLoS pathogens 14, e1007058 (2018).

22. Magalhães, A. C. et al. Peroxisomes are platforms for cytomegalovirus’ evasion from the cellular immune response. Scientific reports 6, 26028 (2016).

23. Zheng, C. & Su, C. Herpes simplex virus 1 infection dampens the immediate early antiviral innate immunity signaling from peroxisomes by tegument protein VP16. Virology journal 14, 35 (2017).

24. Barton, E., Mandal, P. & Speck, S. H. Pathogenesis and host control of gammaherpesviruses: lessons from the mouse. Annual review of immunology 29, 351–397 (2011).

25. Reese, T. et al. Helminth infection reactivates latent γ-herpesvirus via cytokine competition at a viral promoter. Science 345, 573–577 (2014).

26. Reese, T. A. Co-infections: Another Variable in the Herpesvirus Latency-Reactivation Dynamic. Journal of virology 90, JVI.01865-15-5537 (2016).

27. Reddy, J. & Hashimoto, T. Peroxisomal beta-oxidation and peroxisome proliferator-activated receptor alpha: an adaptive metabolic system. Annual review of nutrition 21, 193–230 (2001).

28. Teissier, E. et al. Peroxisome proliferator-activated receptor alpha induces NADPH oxidase activity in macrophages, leading to the generation of LDL with PPAR-alpha activation properties. Circulation research 95, 1174–82 (2004).

29. Nguyen, T., Nioi, P. & Pickett, C. B. The Nrf2-antioxidant response element signaling pathway and its activation by oxidative stress. The Journal of biological chemistry 284, 13291–5 (2009).

30. Chan, K., Lu, R., Chang, J. & Kan, Y. NRF2, a member of the NFE2 family of transcription factors, is not essential for murine erythropoiesis, growth, and development. Proceedings of the National Academy of Sciences of the United States of America 93, 13943–8 (1996).

31. Itoh, K. et al. An Nrf2/small Maf heterodimer mediates the induction of phase II detoxifying enzyme genes through antioxidant response elements. Biochemical and biophysical research communications 236, 313–22 (1997).

32. McMahon, M. et al. The Cap’n’Collar basic leucine zipper transcription factor Nrf2 (NF-E2 p45-related factor 2) controls both constitutive and inducible expression of intestinal detoxification and glutathione biosynthetic enzymes. Cancer research 61, 3299–307 (2001).

33. Dutia, B., Allen, D., Dyson, H. & Nash, A. Type I interferons and IRF-1 play a critical role in the control of a gammaherpesvirus infection. Virology 261, 173–179

34. Hwang, S. et al. Persistent gammaherpesvirus replication and dynamic interaction with the host in vivo. Journal of virology 82, 12498–12509 (2008).

35. den Bossche, J., O’Neill, L. A. & Menon, D. Macrophage Immunometabolism: Where Are We (Going)? Trends in immunology 38, 395–406 (2017).

36. Hall, C. J. et al. Immunoresponsive gene 1 augments bactericidal activity of macrophage-lineage cells by regulating β-oxidation-dependent mitochondrial ROS production. Cell metabolism 18, 265–78 (2013).

37. Moon, J.-S. S. et al. NOX4-dependent fatty acid oxidation promotes NLRP3 inflammasome activation in macrophages. Nature medicine 22, 1002–12 (2016).

38. Bulua, A. C. et al. Mitochondrial reactive oxygen species promote production of proinflammatory cytokines and are elevated in TNFR1-associated periodic syndrome (TRAPS). The Journal of experimental medicine 208, 519–33 (2011).

39. Dixit, E. et al. Peroxisomes are signaling platforms for antiviral innate immunity. Cell 141, 668–81 (2010).

40. Odendall, C. et al. Diverse intracellular pathogens activate type III interferon expression from peroxisomes. Nature immunology 15, 717–26 (2014).

41. Shang, G., Zhang, C., Chen, Z. J., Bai, X.-C. C. & Zhang, X. Cryo-EM structures of STING reveal its mechanism of activation by cyclic GMP-AMP. Nature 567, 389–393 (2019).

42. Ergun, S. L., Fernandez, D., Weiss, T. M. & Li, L. STING Polymer Structure Reveals Mechanisms for Activation, Hyperactivation, and Inhibition. Cell 178, 290–301.e10 (2019).

43. Jin, L., Lenz, L. L. & Cambier, J. C. Cellular reactive oxygen species inhibit MPYS induction of IFNβ. PloS one 5, e15142 (2010).

## REFERENCES

1. Reese, T. et al. Helminth infection reactivates latent γ-herpesvirus via cytokine competition at a viral promoter. Science 345, 573–577 (2014).

2. Weck, K. E. et al. Murine γ-herpesvirus 68 causes severe large-vessel arteritis in mice lacking interferon-γ responsiveness: A new model for virus-induced vascular disease. Nature medicine 3, 1346–1353 (1997).

3. Hwang, S. et al. Persistent gammaherpesvirus replication and dynamic interaction with the host in vivo. Journal of virology 82, 12498–12509 (2008).

4. Rocke, D. & Durbin, B. A model for measurement error for gene expression arrays. Journal of computational biology: a journal of computational molecular cell biology 8, 557–69 (2001).

5. Dozmorov, I. & Lefkovits, I. Internal standard-based analysis of microarray data. Part 1: analysis of differential gene expressions. Nucleic acids research 37, 6323–39 (2009).

6. Dozmorov, I. & Centola, M. An associative analysis of gene expression array data. Bioinformatics (Oxford, England) 19, 204–11 (2003).

